# OPSTA: An Online Analysis Platform for Chinese Sports Science Thesis Data Based on the Shiny Framework

**DOI:** 10.64898/2025.12.26.696552

**Authors:** Lidian Meng, He Zheng

## Abstract

With the continuous expansion of postgraduate enrollment in sports science in China, problems related to research design and statistical analysis in master’s theses have become increasingly prominent. Some postgraduate students have limited statistical training, which may lead to inappropriate research designs, improper selection of statistical methods, failure to check underlying assumptions, and insufficient reporting of effect sizes and confidence intervals, ultimately undermining the scientific rigor and credibility of research conclusions. To address these issues systematically, it is necessary to develop an easy-to-use data analysis tool tailored to the sports science domain and aligned with statistical reporting standards. Accordingly, this study developed the Online Platform for Sports Thesis Analysis (OPSTA). Built on the Shiny framework, OPSTA integrates statistical procedures commonly used in sports science research, including descriptive statistics, independent-samples and paired-samples t tests, one-way and multifactor analysis of variance (ANOVA), correlation analysis, regression analysis, and reliability and validity testing. Through an interactive and visualized interface, the platform guides users through data import, method selection, and result export. While lowering the barrier to statistical analysis, OPSTA emphasizes assumption checking and standardized result presentation, thereby reducing the misuse of statistical methods and improving reporting quality. The platform is available at: https://menglab.org.cn/opsta/.

## 1 Introduction

In recent years, with the continuous expansion of postgraduate education in China, enrollment in master’s programs in sports science has increased rapidly, and the training system has gradually shifted from an elite-oriented model to a more inclusive one. Against this background, a growing mismatch has emerged between the limited statistical foundations and insufficient research training of some postgraduate students and the research design and data analysis competencies required for high-quality master’s theses. Deficiencies in problem formulation, experimental or survey design, sample size estimation, and variable control are frequently observed in theses, directly undermining the scientific rigor and credibility of research conclusions and posing new challenges to the quality of sports science master’s theses.

On the one hand, some postgraduate students in sports science mechanically apply parametric statistical methods—such as *t* tests, analysis of variance, correlation analysis, and regression analysis—without examining underlying statistical assumptions, while neglecting the reporting of effect sizes and confidence intervals. This practice often leads to the misinterpretation of *P* values and, in some cases, the emergence of spurious statistically significant findings. On the other hand, questionable research practices, including multiple testing, selective reporting, arbitrary handling of outliers, and even data fabrication or data trading, have to some extent exacerbated the dissemination of false research findings. These behaviors undermine the credibility and reproducibility of sports science research and may mislead subsequent studies and practical applications.

In response to these issues, it is necessary to systematically examine common errors in research design and statistical analysis in sports science master’s theses and to develop a dedicated statistical analysis tool tailored to the field. Accordingly, this study developed the Online Platform for Sports Thesis Analysis (OPSTA). The platform integrates commonly used statistical functions, including descriptive statistics, *t* tests, analysis of variance, correlation and regression analyses, as well as reliability and validity testing. Through a visualized and interactive interface, OPSTA lowers the barrier to statistical analysis and provides sports science postgraduate students with a tool that balances methodological rigor and operational usability. By objectively identifying existing problems, the platform aims to offer a feasible pathway for improving statistical practice and enhancing the overall quality of sports science research and master’s theses.

## 2 Methods

### 2.1 Development of the OPSTA Online Analysis Platform

#### 2.1.1 Platform Architecture

OPSTA is an online statistical analysis platform developed based on R version 4.5.1 and the R Shiny framework (version 1.11.1). All components of the platform are currently deployed on the JD Cloud North China data center. The front-end interface and statistical analysis functions are implemented through the coordinated use of multiple R extension packages. Specifically, (1) the overall interface theme is configured using shinythemes (version 1.2); (2) data preprocessing, missing value handling, and descriptive statistics are primarily implemented using Hmisc (version 5.2.4), plyr (version 1.8.9), and Rmisc (version 1.5.1); (3) graphical visualization is constructed using ggplot2 (version 4.0.1), supplemented by qqplotr (version 0.0.7) and ggpubr (version 0.6.2) for the generation of normality assessment plots and related statistical graphics; (4) scale reliability and validity analyses are conducted using psych (version 2.5.6); (5) regression models are fitted using built-in R functions, followed by further diagnostics—such as analysis of covariance, multicollinearity assessment, and model checking—performed with the car package (version 3.1.3); (6) repeated-measures data are analyzed by fitting linear mixed-effects models using nlme (version 3.1.168); and (7) final analytical results are presented as interactive data tables via DT (version 0.34), facilitating result filtering, sorting, and export.

#### 2.1.2 Platform Workflow

The overall workflow of OPSTA consists of three steps (Figure 1). First, users upload their data by saving it in comma-separated values (.csv) format and importing it into the platform; standardized example files are provided for reference. Second, users select appropriate statistical analysis modules according to their research objectives to process and analyze the data. Third, the analytical results can be directly copied for use, and the generated figures can be saved and exported for reporting and manuscript preparation.

**Figure 1.**
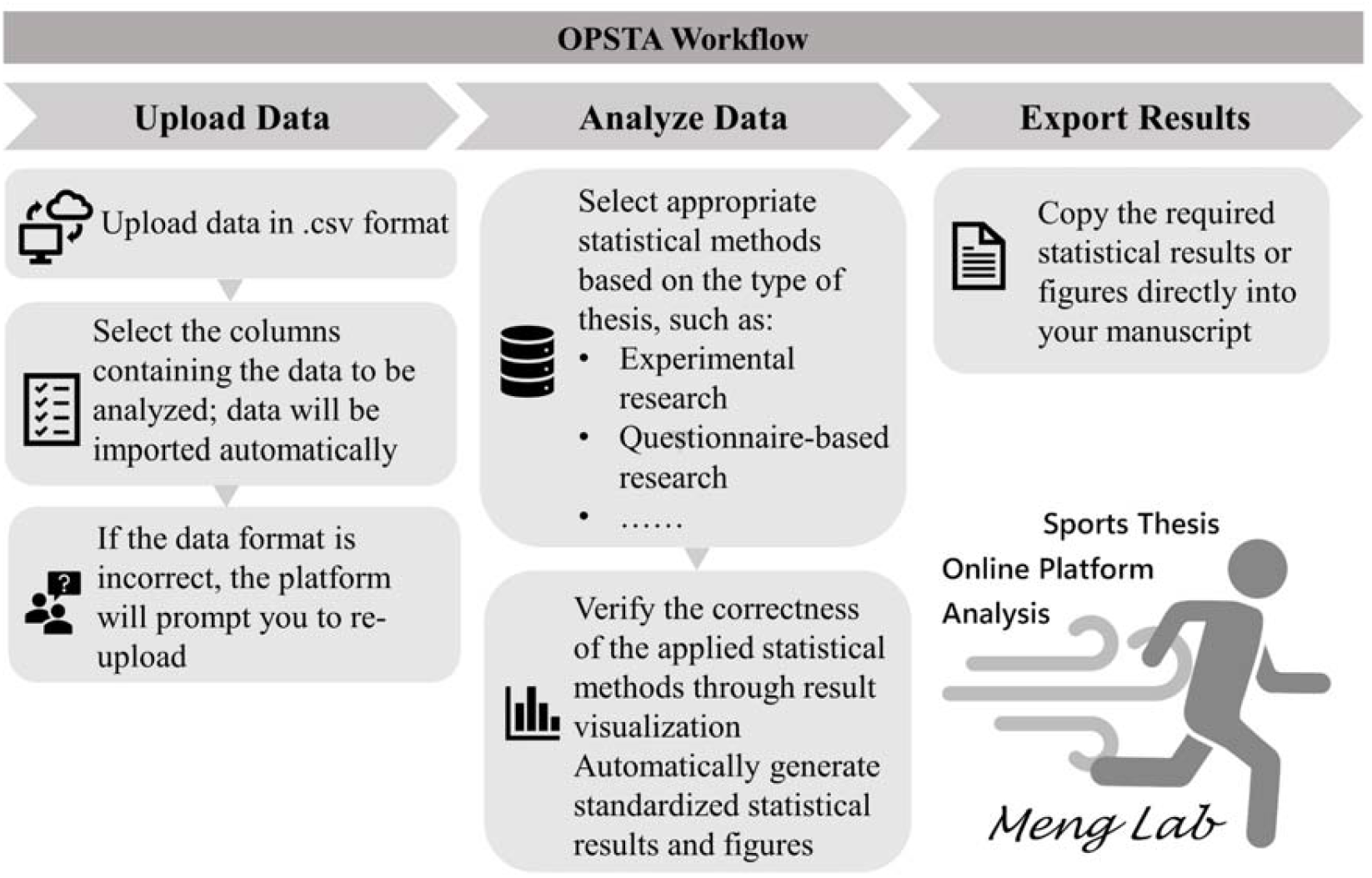
Workflow of the OPSTA Platform

### 2.2 Visualized Application of OPSTA

To demonstrate the practical application of OPSTA, this study employed a human standing long jump dataset released by Yu Zhangguo and colleagues from Beijing Institute of Technology. The dataset was collected using the Xsens MVN Link motion capture system and includes multidimensional kinematic parameters recorded at major body segments during the standing long jump task. These parameters comprise quaternions, Euler angles, position, velocity, acceleration, angular velocity, and angular acceleration. The recorded body segments include the trunk (pelvis, L3 and L5 lumbar vertebrae, T8 and T12 thoracic vertebrae, neck, and head), the left lower limb (thigh, shank, foot, and toe), the right lower limb (thigh, shank, foot, and toe), the left upper limb (upper arm, forearm, and hand), and the right upper limb (upper arm, forearm, and hand).

As an illustrative example, this study focused on the three-dimensional pelvic acceleration (m/s^2^) across different time frames during the standing long jump. The dataset contains 2,056 observations and 70 variables. The analyses presented are intended solely to demonstrate the operational procedures and analytical capabilities of the platform and do not carry substantive research implications. Use of the dataset was approved by the National Basic Science Data Center on July 3, 2025 (approval number: CSTR:16666.11.nbsdc.pqyexbfy).

## 3 Results

### 3.1 Overview of the OPSTA Platform

The OPSTA platform features a concise and intuitive interface with a modular design, allowing users to rapidly access different statistical analysis modules via the top navigation bar and thereby reducing the operational complexity of statistical analyses (Figure 2). The left panel of the main interface presents the platform name, functional positioning, and development team information, clearly indicating its application scenario for sports science thesis data analysis. The right panel serves as the core functional area and is organized according to common data analysis workflows in sports science research. It includes modules such as “About the Platform,” “Upload Data,” “Bivariate Analysis,” “Multivariate Comparison,” “Categorical Tests,” “Reliability and Validity Analysis,” “*t* Tests,” and “One-Way and Multifactor Analysis of Variance.”

**Figure 2.**
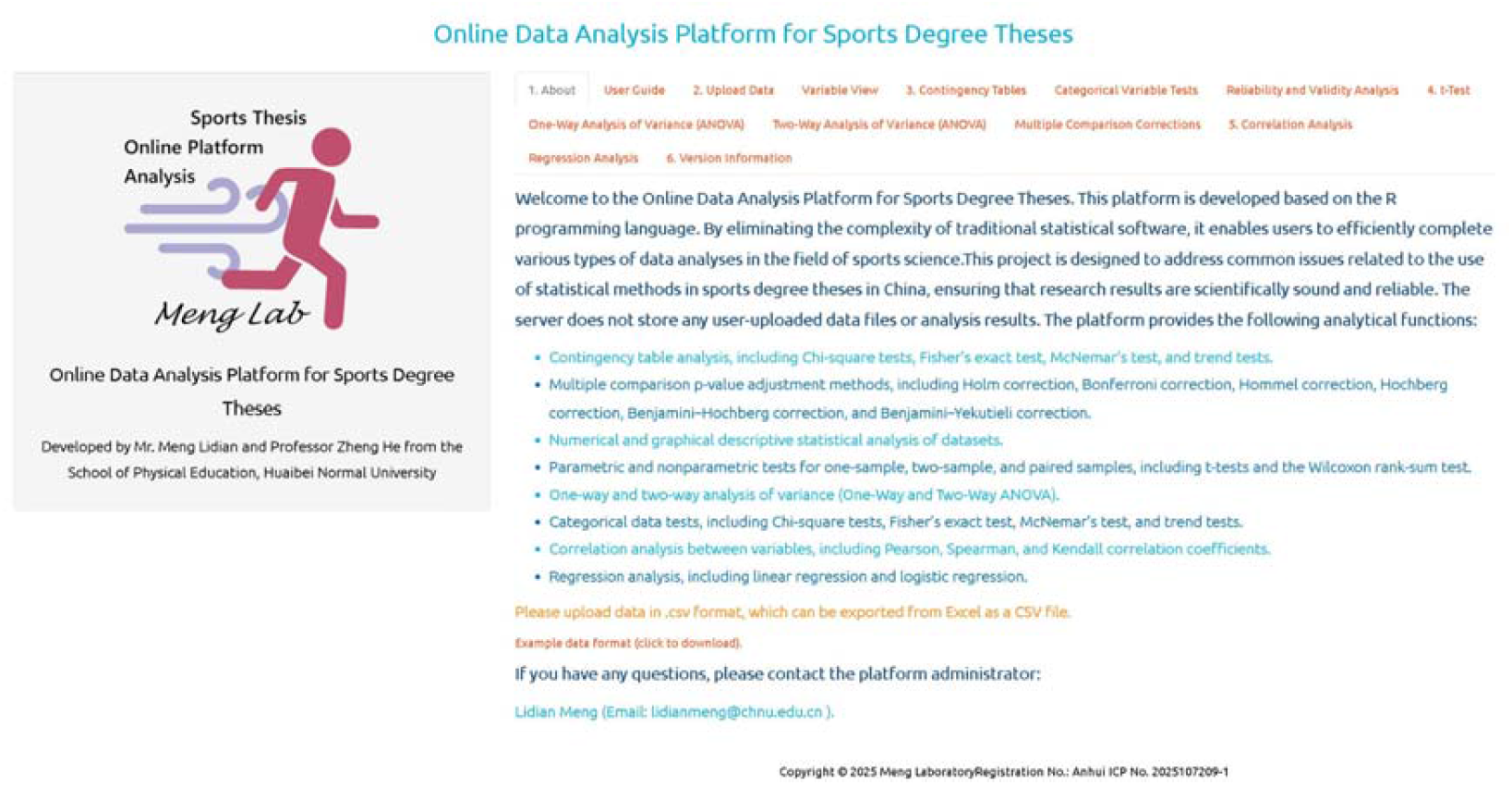
Main interface of the OPSTA platform

In terms of data handling, the platform supports the upload of CSV-formatted data files and provides standardized example datasets to help novice users understand data structure requirements. The statistical analysis functions cover methods commonly used in sports science research, including chi-square tests, Fisher’s exact test, McNemar’s test, trend tests, multiple comparison correction procedures (e.g., Holm, Bonferroni, Hochberg, and Benjamini–Hochberg), as well as Pearson, Spearman, and Kendall correlation analyses, linear regression, and logistic regression analyses.

With respect to result presentation, OPSTA automatically outputs corresponding statistical results and visualizations, facilitating intuitive interpretation of analytical outcomes. The platform places particular emphasis on assumption checking and standardized result reporting, which helps reduce the misuse of statistical methods and nonstandard reporting practices in sports science master’s theses.

### 3.2 Example of Visualized Application Using OPSTA

Changes in pelvic acceleration along the X, Y, and Z axes were used as an illustrative example (Figure 3). Shapiro–Wilk normality tests indicated that acceleration data in all three directions deviated from a normal distribution (X: *W* = 0.355, *P* < 0.001; Y: *W* = 0.383, *P* < 0.001; Z: *W* = 0.528, *P* < 0.001). Accordingly, Wilcoxon paired-rank tests were applied to compare differences among the three directions at the same sampling points. The median difference between X and Y was not statistically significant (*P* = 0.476), whereas statistically significant differences were observed between X and Z (*P* < 0.001) and between Y and Z (*P* = 0.027).

**Figure 3.**
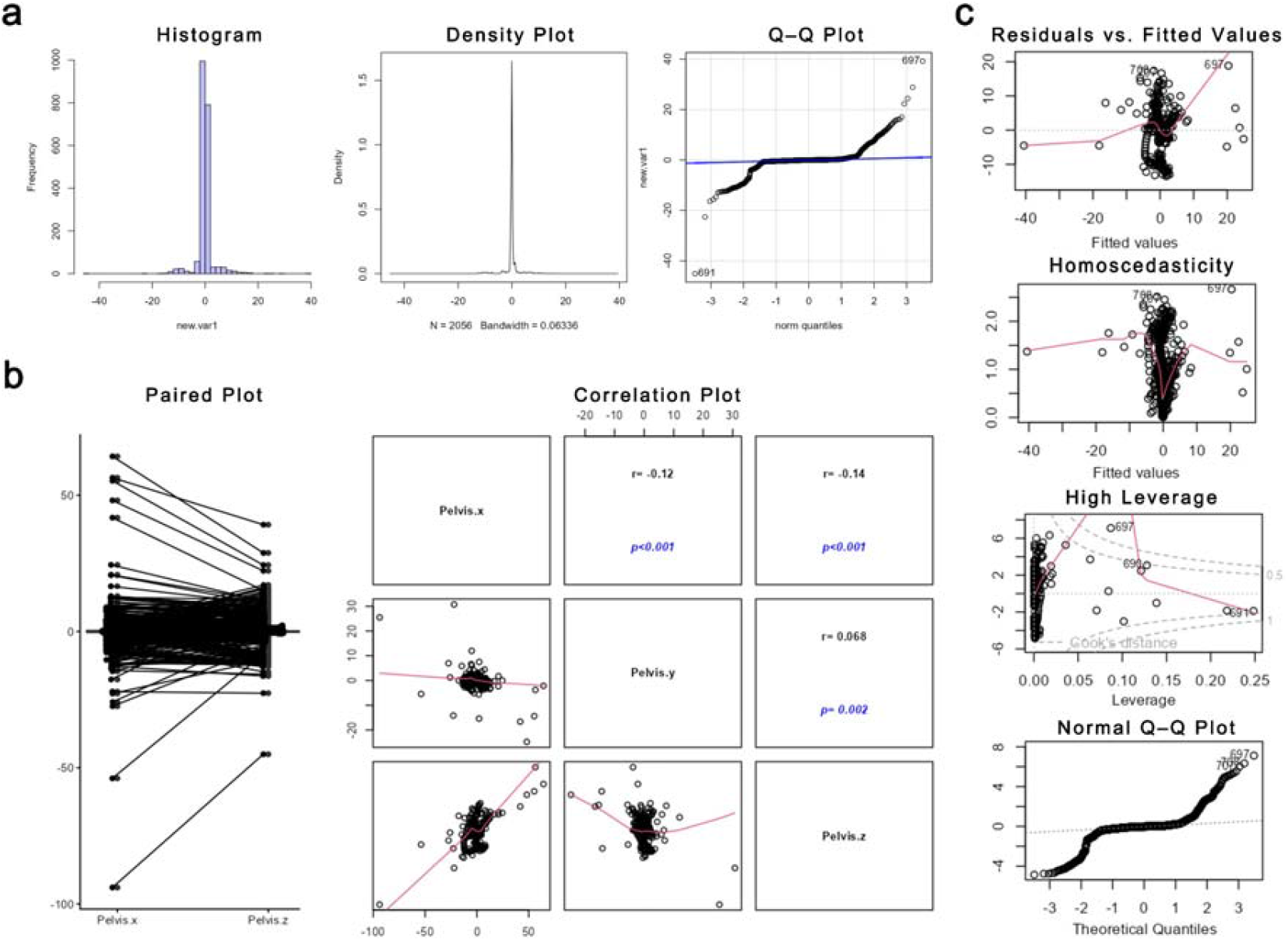
Statistical analysis and visualization of pelvic three-dimensional acceleration data using OPSTA. (a) Histogram, kernel density curve, and Q–Q plot of pelvic acceleration in the Z direction, illustrating data distribution characteristics; (b) Paired plot of X and Z, as well as scatterplots and Spearman correlation coefficients for pairwise comparisons among X, Y, and Z directions, illustrating directional differences and associations; (c) Diagnostic plots of the multiple linear regression model, including residuals versus fitted values, scale–location plot, leverage and influence plots, and residual normal Q–Q plot, used to assess linearity, homoscedasticity, outliers, and residual normality assumptions.

Spearman rank correlation analysis was further conducted to assess associations among the X, Y, and Z directions. Very weak negative correlations were observed between X and Y (*r* = −0.122, *P* < 0.001) and between X and Z (*r* = −0.136, *P* < 0.001), while a very weak positive correlation was found between Y and Z (*r* = 0.068, *P* = 0.002). Overall, the magnitudes of these correlations were low.

Subsequently, a multiple linear regression model was constructed with Z-direction acceleration as the dependent variable and X- and Y-direction accelerations as independent variables. The overall model was statistically significant (*F* = 436.3, *P* < 0.001), with X and Y jointly explaining approximately 29.8% of the variance in Z (*R*^2^ = 0.298). After controlling for Y, X showed a significant positive predictive effect on Z (β = 0.337, *P* < 0.001), whereas after controlling for X, Y exhibited a significant negative predictive effect on Z (β = −0.349, *P* < 0.001).

## 4 Discussion

In research fields such as biology, medicine, and psychology, the platformization and automation of research workflows have been widely adopted, leading to the development of a series of online analytical platforms represented by tools such as Galaxy and ExPASy. These platforms integrate data analysis, result visualization, and quality control within a unified framework, thereby enhancing the transparency, standardization, and reproducibility of research processes. Through modularized design, platform-based workflows also reduce manual intervention, which can effectively mitigate analytical bias arising from insufficient researcher experience or improper operations. As the volume and structural complexity of research data continue to increase, online analytical platforms that require no local software installation have become an important means of improving both research efficiency and research quality.

By contrast, statistical practices in sports science research have long relied heavily on individual researcher experience, resulting in substantial variability in the selection of statistical methods and a lack of unified standards for result reporting. The issues observed in this study—including the misuse of statistical methods, insufficient reporting of effect sizes, and inconsistent use of statistical terminology—indicate that statistical practice in sports science remains relatively fragmented. Many researchers are not proficient in standardized programming languages and instead depend primarily on complex menu-driven operations in statistical software. This approach increases arbitrariness in method selection and makes it difficult to ensure consistency and 规范性 in reporting. Such a highly experience-dependent analytical paradigm renders research quality overly sensitive to individual judgment, thereby weakening the robustness and reproducibility of research findings.

Developing an online statistical analysis platform tailored to the characteristics of sports science research can help integrate statistical processing into a unified analytical framework. Such platforms can improve reporting quality through standardized output templates, reduce the likelihood of statistical misuse through built-in assumption checks and quality prompts, and enhance reproducibility across studies. Moreover, platform-based analytical tools hold substantial value for teaching and research training, as they enable researchers to gradually develop rigorous and standardized statistical thinking through hands-on use. Consequently, in the context of expanding sports research, increasing interdisciplinary collaboration, and growing data complexity, the construction and dissemination of online analytical platforms are not only of immediate practical significance but also of long-term value for advancing the methodological development of sports science.

